# Lateralized phenotypic differences after intrahippocampal kainic acid injection in female mice

**DOI:** 10.1101/2021.09.09.459684

**Authors:** Cathryn A. Cutia, Leanna K. Leverton, Xiyu Ge, Rana Youssef, Lori T. Raetzman, Catherine A. Christian-Hinman

**Author notes:** Corresponding author: Catherine A. Christian-Hinman, Ph.D., 407 S. Goodwin Ave., Urbana, IL 61801 USA. Equal contribution (Cathryn A. Cutia, Leanna K. Leverton).

## Abstract

Clinical evidence indicates that patients with temporal lobe epilepsy (TLE) often show differential outcomes of comorbid conditions in relation to the lateralization of the seizure focus. A particularly strong relationship exists between the side of seizure focus and the propensity for distinct reproductive endocrine comorbidities in women with TLE. Therefore, here we evaluated whether targeting of left or right dorsal hippocampus for intrahippocampal kainic acid (IHKA) injection, a model of TLE, produces different outcomes in hippocampal granule cell dispersion, body weight gain, and multiple measures of reproductive endocrine dysfunction in female mice. One, two, and four months after IHKA or saline injection, *in vivo* measurements of estrous cycles and weight were followed by *ex vivo* examination of hippocampal dentate granule cell dispersion, circulating ovarian hormone and corticosterone levels, ovarian morphology, and pituitary gene expression. IHKA mice with right-targeted injection (IHKA-R) showed greater granule cell dispersion and pituitary *Fshb* expression compared to mice with left-targeted injection (IHKA-L). By contrast, pituitary expression of *Lhb* and *Gnrhr* were higher in IHKA-L mice compared to IHKA-R, but these values were not different from respective saline-injected controls. IHKA-L mice also showed an increased rate of weight gain compared to IHKA-R mice. Increases in estrous cycle length, however, were similar in both IHKA-L and IHKA-R mice. These findings indicate that although major reproductive endocrine dysfunction phenotypes present similarly after targeting left or right dorsal hippocampus for IHKA injection, distinct underlying mechanisms based on lateralization of epileptogenic insult may contribute to produce similar emergent reproductive endocrine outcomes.

**Highlights:**

- Greater granule cell dispersion with right-sided IHKA injection
- Changes in pituitary gene expression vary with side of injection
- Increased weight gain after left-sided injection
- Similar estrous cycle disruption after injection of left or right hippocampus

## 1. Introduction

Temporal lobe epilepsy (TLE) is the most prevalent form of focal epilepsy in adults (Engel, 2001). Patients with TLE commonly develop comorbid conditions such as cognitive impairment (Helmstaedter & Kockelmann, 2006), psychopathologies (Kandratavicius et al., 2012), and reproductive endocrine dysfunction (Herzog et al., 1986b, 1986a). Such comorbidities likely occur as results of a number of structural and functional abnormalities that develop both within the temporal lobe (Berkovic et al., 1991; Blümcke et al., 2002; Sutula et al., 1989) and elsewhere in the brain (Gross et al., 2006; Keller & Roberts, 2008; Campos et al., 2016). As the human brain shows a high level of structural and functional lateralization (Gazzaniga, 1995), such abnormalities can manifest differently based on the hemisphere in which the seizure focus, the site of seizure origination, resides. For example, patients with left-sided seizure foci often display more widespread abnormalities affecting the contralateral hemisphere than patients with right-sided seizure foci (Campos et al., 2016; Coan et al., 2009; Haneef et al., 2014; Keller et al., 2012). Furthermore, there is some evidence to suggest that cognitive impairment shows lateralization based on seizure focus, as patients with left-sided seizure foci exhibit more cognitive impairment than those with a right-sided focus (Alessio et al., 2006; Bonilha et al., 2007; Phuong et al., 2021).

Particularly prominent indications for differential comorbid outcomes based on seizure focus lateralization are the clinical findings of distinct forms of reproductive endocrine dysfunction among women with TLE. Specifically, women with left temporal lobe seizure foci have increased rates of developing polycystic ovary syndrome (PCOS), whereas those with right-sided foci exhibit higher rates of hypothalamic amenorrhea and hypogonadotropic hypogonadism (Herzog, 1993; Herzog et al., 1986b; Kalinin & Zheleznova, 2007). Therefore, it appears that the laterality of the seizure focus could be a significant risk factor for particular reproductive endocrine disorders or other outcomes, but the mechanisms producing this relationship remain unknown. Furthermore, this correlation has been difficult to replicate in animal models of TLE. For example, one study in rats found no difference in impacts on the estrous cycle after kindling the left or right amygdala (Hum et al., 2009). To date, no animal model of chronic epilepsy with recurrent spontaneous seizures has definitively shown differential effects of seizure focus lateralization on endocrine function or other phenotypic outcomes. Characterization of animal models investigating the impacts of left- vs. right-sided seizure foci could enable the study of mechanisms linking lateralization of epileptic focus to comorbid reproductive endocrine disorders and other distinct structural or functional changes. Furthermore, such an investigation would indicate whether lateralization of the epileptogenic insult contributes to differential experimental outcomes.

Several rodent models of epilepsy exhibit reproductive endocrine dysfunction in the form of disrupted estrous cycles in females (Amado et al., 1993; Christian et al., 2020; Edwards et al., 1999; Scharfman et al., 2008) and thus could potentially be used to study distinct outcomes in association with seizure focus lateralization. Of these models, the intrahippocampal kainic acid (IHKA) mouse model not only leads to disruption in cycles (Li et al., 2017), but also shows similar histological and seizure patterning characteristics as human TLE (Bouilleret et al., 1999; Gröticke et al., 2008; Riban et al., 2002). This model also allows for the epileptogenic insult to be discretely targeted to the left or right hemisphere to develop a specific, lateralized seizure focus similar to those often seen in patients with TLE.

The pituitary gonadotropins, luteinizing hormone (LH) and follicle-stimulating hormone (FSH), are largely responsible for gonadal steroidogenesis and ovarian follicle development necessary for normal reproduction. The expression of gonadotropins at the pituitary is initiated when gonadotropin-releasing hormone (GnRH) binds its receptor, GnRH-R, at the pituitary. Elucidating whether expression levels of the functional subunits of these hormones and the GnRH-R is altered at the pituitary level is vital to understand input to the system and gives basic insight into the functional output. However, changes in pituitary gene expression have not been studied in IHKA animals.

Here, the phenotypic effects of left-versus right-sided dorsal IHKA injection were compared in adult female mice to test the hypothesis that differential outcomes are produced based on the side targeted for injection. This study focused on female mice as it is largely women rather than men with TLE that demonstrate evidence for differential reproductive endocrine outcomes based on laterality of the seizure focus (Herzog, 1993; Kalinin & Zheleznova, 2007). Hippocampal granule cell dispersion and gliosis, pituitary gonadotropin gene expression levels, estrous cycle lengths, ovarian histology, gonadal and stress hormone levels, and post-injection weight gain were assessed across several time points after injection.

## 2. Materials and Methods

### 2.1 Animals and estrous cycle monitoring

All animal procedures complied with the ARRIVE guidelines and were approved by the Institutional Animal Care and Use Committee of the University of Illinois Urbana-Champaign. Female C57BL/6J mice (#000664, Jackson Laboratories) were purchased for delivery at 6 weeks of age and housed in a 14:10 h light:dark cycle with food and water available *ad libitum*. Starting 5 days after arrival, daily estrous cycle monitoring was performed between 0900 to 1100 h (relative to 1900 h lights off time) using a vaginal cytology protocol described previously (Pantier et al., 2019). Mice were assessed for a minimum of 14 days to verify regular cycling prior to saline/KA injection as described below. As outlined in **Figure 1**, the mice followed a timeline of estrous cycling that resumed beginning 3 weeks after injection for a minimum of 14 days, after which a subset of mice were euthanized by decapitation for tissue collection (1-month groups). At 7 weeks post-injection, the remaining mice resumed cycle monitoring for a minimum of 14 days, and a second subset was collected (2-months groups). At 15 weeks post-injection, the remaining mice resumed cycle monitoring again, and samples were collected for the 4-months groups. Cycle length (period) was calculated as the number of days required to progress from one day of estrus through each stage to the next day of estrus.

**Figure 1.**
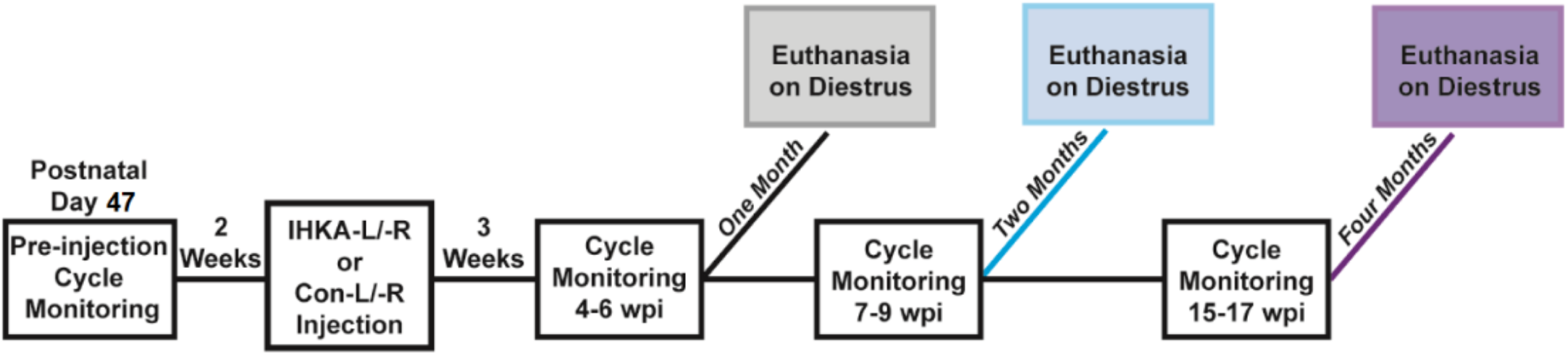
Experimental timeline. Schematic of procedures and timepoints for each component. Note that each timepoint has 4 groups: IHKA-L, IHKA-R, Con-L, and Con-R. At the time of euthanasia, trunk blood, pituitary, brain, and ovaries were collected from each animal. wpi = weeks post-injection.

### 2.2 IHKA injection surgeries and post-operative acute seizure video monitoring

Mice aged postnatal day 60 and older underwent stereotaxic unilateral injection of KA (Tocris Bioscience; 50 nL of 20 mM prepared in 0.9% sterile saline) into the left or right dorsal hippocampal region (from Bregma: 2.0 mm posterior, 1.5 mm lateral, 1.4 mm ventral to the cortical surface) as previously described (Li et al., 2017). All injections were performed on diestrus. Age-matched controls were injected with the same volume of saline at the same location. All surgeries were performed under isoflurane anesthesia (2-3%, vaporized in 100% oxygen). During recovery immediately after injection, IHKA mice were video recorded (iSpyConnect) following injection via webcam (Logitech C930e). Videos were scored using the Racine scale (Racine, 1972), and the time and stage of the first behavioral seizure at or above stage 3 were recorded.

### 2.3 Tissue collection

All tissue collections were performed on diestrus between 1200 and 1600 h. Mouse weights were recorded prior to euthanasia by decapitation. The trunk blood, brain, pituitary gland, and ovaries were collected. Trunk blood was stored at 4°C for 24 h, then spun in a centrifuge for 10-15 minutes at 4°C. Serum was then isolated and stored at −80°C. Brains were bulk-fixed at 4°C in 4% paraformaldehyde (Sigma-Aldrich 158127) for 24 h, then switched to 30% sucrose with 5% sodium azide (ACROS Organics 19038-1000) and stored at 4°C. Pituitaries were stored in RNAlater Stabilization Solution (Invitrogen, AM7020) at −20°C. Ovaries were stored in 10% formalin at 4°C overnight, then switched to 100% ethanol and stored at 4°C.

### 2.4 Hippocampal granule cell dispersion quantification

40-μm coronal hippocampal sections were collected using a freezing microtome (SM 2010R, Leica Biosystems). Four to six sections per mouse from the dorsal hippocampal region were used for evaluation of hippocampal granule cell dispersion by cresyl violet (Nissl) staining and/or gliosis by glial fibrillary acidic protein (GFAP) staining. For cresyl violet staining, sections were mounted on charged glass slides, stained with cresyl violet (Sigma-Aldrich C5042) for 12 minutes at room temperature (∼22°C), dehydrated with graded ethanol solutions (70-100%), cleaned in xylene, and coverslipped with Permount. For GFAP immunofluorescence, sections were incubated with an anti-GFAP mouse monoclonal antibody (1:1000, Sigma-Aldrich G3893) for 48 h at 4°C on a shaker, followed by incubation with DyLight 594 horse anti-mouse secondary (1:1000, Vector DI-2594) for 2 h at room temperature on a shaker. Sections were then mounted on charged glass slides and coverslipped using Vectashield Hardset Antifade Mounting Medium with DAPI (Vector Laboratories H-1500). Image acquisition was performed using an Olympus BX43 brightfield and fluorescence microscope equipped with an Infinity 3-6UR Teledyne Lumenera Camera and Infinity Capture software (Lumenera). The degree of hippocampal damage produced after IHKA injection was quantified by measuring the surface area of the dentate gyrus granule cell layer and hilar regions in ImageJ (Lisgaras & Scharfman, 2021). The surface area of the ipsilateral dentate gyrus and hilar regions was then compared to the contralateral regions in the same mouse. To account for normal variations in granule cell layer area in different regions of the hippocampus, the relative distance from Bregma was attained by comparisons to a mouse brain atlas (Paxinos & Keith B.J. Franklin, 2019) for each hippocampal section. Only ipsilateral and contralateral hippocampal regions with the same relative distance from Bregma were compared to one another to determine the ipsilateral:contralateral ratio. The integrated density of GFAP was analyzed in ImageJ and normalized to the overall size of each ipsilateral hippocampal region, as contralateral hippocampal regions did not exhibit significant amounts of GFAP expression.

### 2.5 Hormone assays

Enzyme-linked immunosorbent assays (ELISAs) for progesterone, estradiol, testosterone, and corticosterone were performed on trunk blood serum samples according to the manufacturers’ instructions (progesterone: IBL America (IB79183); estradiol: Calbiotech (ES180S-100); testosterone: IBL America (IB79174); corticosterone: Arbor Assays (K014-H1)) and evaluated using a Bio-Tek 800TS Microplate Absorbance Reader. As needed, samples were diluted to match the volume required for the testing. Samples were run in duplicate where appropriate and the average of the duplicate was used as the final hormone concentration value for each mouse. Intra-assay coefficient of variation (CV) values were calculated from the standard deviations within plates. Samples with CV values higher than 10 were excluded from the dataset, and plate variation was controlled for as a factor in statistical analyses.

### 2.6 Ovary histology and follicle counts

Both left and right ovaries were paraffin-embedded, and six consecutive sections of 5-µm thickness were collected from each ovary and stained with hematoxylin and eosin. An investigator blinded to treatment group screened the sections to quantify the number of primary, secondary, and antral follicles, and corpora lutea. Follicles were defined as primary, secondary, or antral according to characteristic morphology (Myers et al., 2004). The numbers of each type of ovarian follicle and corpora lutea were counted from both the left and right ovaries of each mouse and averaged to generate the final value for each mouse used in statistical analyses.

### 2.7 Pituitary qPCR

RNA was isolated from pituitaries using RNAqueous-Micro kits (Invitrogen). Total RNA was reverse-transcribed according to the manufacturer’s instructions using the ProtoScript Strand cDNA Synthesis kit (New England Biolabs) as previously described (Nantie et al., 2014). Oligonucleotide primers for *Lhb*, *Fshb* and *Gnrhr* (Life Technologies, **Table 1**) were used to amplify gene-specific transcripts by quantitative PCR (qPCR). The expression levels of genes of interest were normalized to peptidylprolyl isomerase A (*Ppia)* mRNA levels; *Ppia* is a common reference gene used as an internal control. The data were analyzed using the standard comparative cycle threshold value method (2^−ΔΔCt^).

**Table 1.**
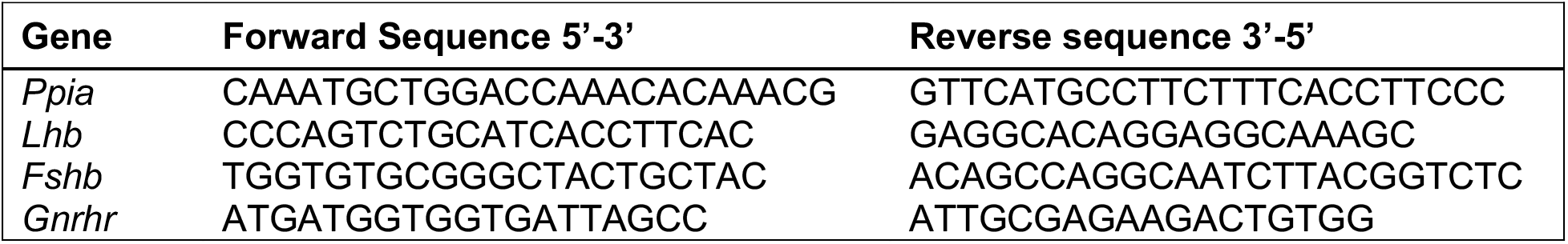
Primer sequences. List of the primer sequences for the gonadotropins (*Lhb, Fshb*), *Gnrhr,* and control *Ppia* used for qPCR.

### 2.8 Statistics

Statistical analyses were conducted in R (R Core Team, 2021). Comparisons between control and IHKA treatment groups and left- vs. right-sided injection groups for hippocampal granule cell dispersion, pituitary mRNA expression levels, and ovary follicle numbers were made using two-way ANOVAs followed by Tukey’s *post hoc* tests. The data for hippocampal granule cell dispersion and pituitary mRNA expression were log-transformed to achieve normality. Comparisions between injection side treatment groups and groups with and without dispersion for GFAP intensity were made using two-way ANOVAs. Comparisons for estrous cycle length were made using two-way ANOVAs (side of injection and saline/KA as factors) and Tukey’s *post hoc* tests for each time point. Comparisons between treatment group, left or right injection group, and time after injection (1, 2, or 4 months) for weight gain were made using three-way ANOVAs and Tukey’s *post hoc* tests. As informed by Box-Cox normality plots, cycle length data were transformed with a negative exponent, and weight data were log-transformed to achieve normality. Comparisons between treatment group, left or right injection group, and ELISA plate group for hormone analyses were made using three-way ANOVAs. Hormone data were log-transformed to achieve normality, which was evaluated with Shapiro-Wilks tests. Correlations between degree of granule cell dispersion and cycle length were made using Pearson’s correlations, and comparisons of proportions of animals exhibiting granule cell dispersion were made with chi-square tests. The criterion for statistical significance was p ≤ 0.05. Results are reported as means ± SEM.

## 3. Results

### 3.1 Evaluation of IHKA injection efficacy

To confirm the efficacy of IHKA injection, all IHKA mice (left-injected = IHKA-L; right-injected = IHKA-R) were screened for behavioral seizures directly following the injection. Some animals could not be recorded for the alotted time due to equipment malfunction. Therefore, all mice that did not show one or more seizure in the recording period, or for which videos were not available, underwent an evaluation for hippocampal sclerosis by Nissl staining. Sections from mice that showed cell death in the CA1 region without significant granule cell layer dispersion underwent further confirmation through immunofluorescent GFAP staining to determine if gliosis was present. If neither granule cell dispersion nor gliosis were observed, and acute seizures could not be confirmed in video screening, the mice were not included in final analyses, in accordance with criteria established previously (Li et al., 2018). A total of 54 IHKA-L mice were established for this study; of these, 12 lacked seizures captured on video immediately following injection. 4 of the 12 animals without video were excluded from the study based on lack of dispersion or gliosis. An additional 3 mice that had one behavioral seizure following injection, but lacked dispersion or gliosis, were also excluded from the study. Of 60 IHKA-R mice entered into the study, 12 lacked videos that captured behavioral seizures following injection. Of these 12 mice, 2 were excluded from the study based on a lack of dispersion or gliosis. An additional 2 mice that had one behavioral seizure following injection, but lacked dispersion or gliosis, were excluded from the study. Therefore, 47 out of 54 IHKA-L mice (87%) and 56 of 60 IHKA-R mice (93%) displayed displayed acute behavioral seizures and/or granule cell dispersion or gliosis. The proportions of mice that displayed seizures and/or showed histopathologies common to IHKA injection for both injection groups (103 of 114 total mice) were thus similar to the ∼90% of mice that are observed to develop chronic recurrent seizures in this model (Rattka et al., 2013).

### 3.2 IHKA-R animals have higher granular cell layer dispersion

To determine the impact of differentially targeting left or right dorsal IHKA injection on hippocampal granule cell dispersion, an additional subset of IHKA animals that met the seizure screening criterion and showed signs of dispersion and/or gliosis, as well as randomly selected saline-injected controls (18 Con-L and 18 Con-R), were added to the histology dataset (**Figure 2A-B**). No saline-injected animals showed signs of cell death, dispersion, or significant levels of GFAP staining. In the resulting analysis of ipsilateral:contralateral (I:C) ratio of the surface area of the dentate gyrus through Nissl staining, there were higher levels of dispersion than in controls (F(1,85) = 13.81, p = 0.0004) in both IHKA-L (p = 0.005) and IHKA-R groups (p < 0.0001). Moreover, side of injection yielded differences (F(1,85) = 6.40, p = 0.01) as IHKA-R animals showed larger dispersion of the granule cell layer compared to IHKA-L (p = 0.04), with no effect of time after injection (F(2,77) = 1.82, p = 0.17, **Figure 2C-D**). From the distribution of I:C ratios, (**Figure 3A**) a threshold ratio of 1.5 was determined to delineate between groups with and without dispersion. The proportion of mice that displayed prominent dispersion (defined as I:C > 1.5) was not different between IHKA-L and IHKA-R groups (χ^2^ = 2.49, p = 0.11). Overall, these data are the first demonstration of differential hippocampal damage based on the side of IHKA injection.

**Figure 2.**
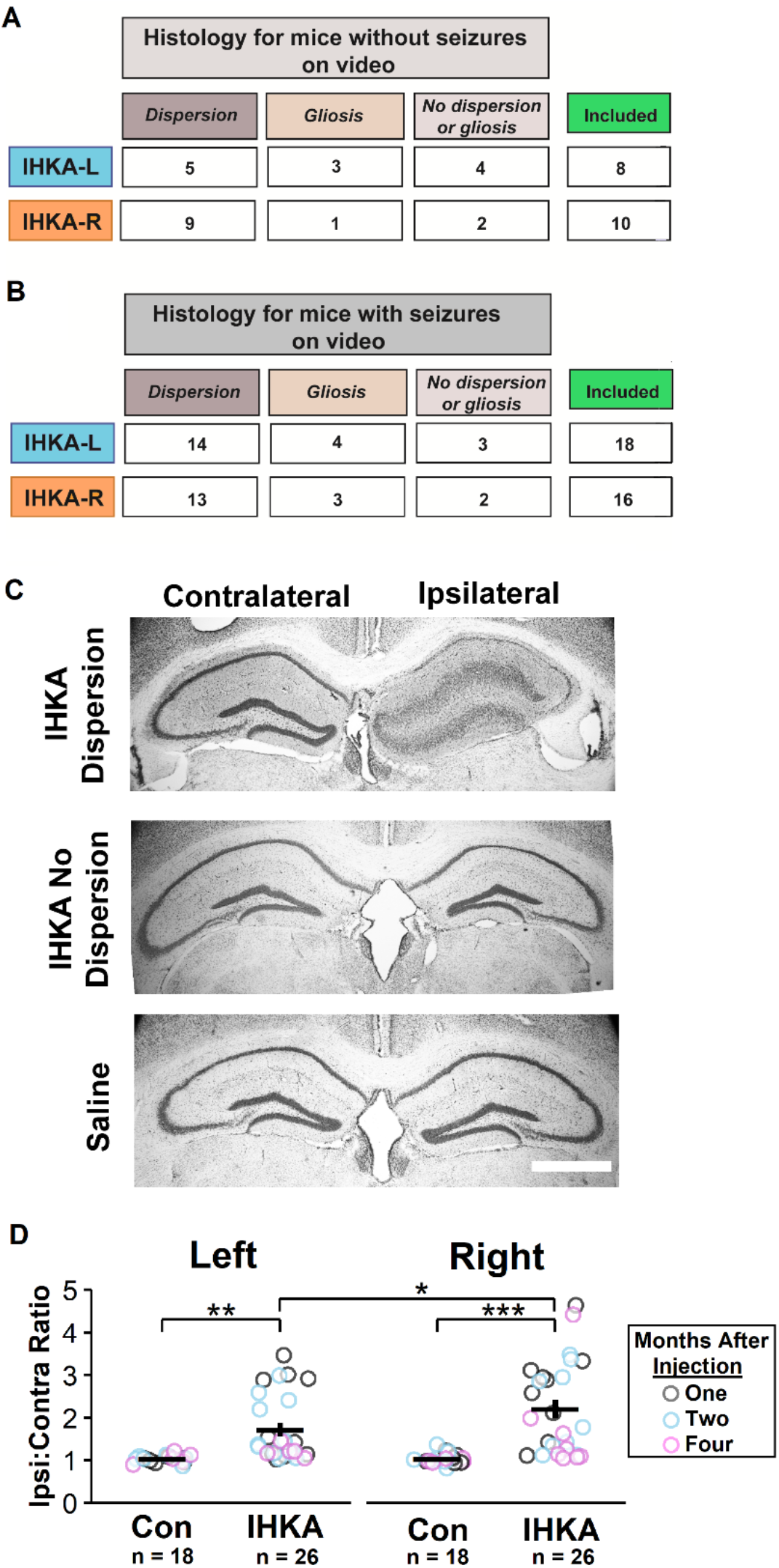
Higher degee of hippocampal granule cell dispersion in IHKA-R compared with IHKA-L mice. (A) Histology inclusion numbers and criteria based on Nissl and GFAP staining for mice that did not show seizures immediately following injection or for which videos were not available. (B) Histology inclusion numbers and criteria based on Nissl and GFAP staining for a randomly selected sample of mice that did show at least one seizure on video collected for 3-5 hours after injection. (C) Representative images of Nissl staining. Scale bar = 1 mm. (D) Individual ipsilateral:contralateral (I:C) ratio of each mouse with mean ± SEM for each treatment group. *p < 0.05, **p<0.01, ***p < 0.001 for comparisons between injection type (Con or IHKA) and injection side (left or right) by two-way ANOVA and Tukey’s *post hoc* tests.

**Figure 3.**
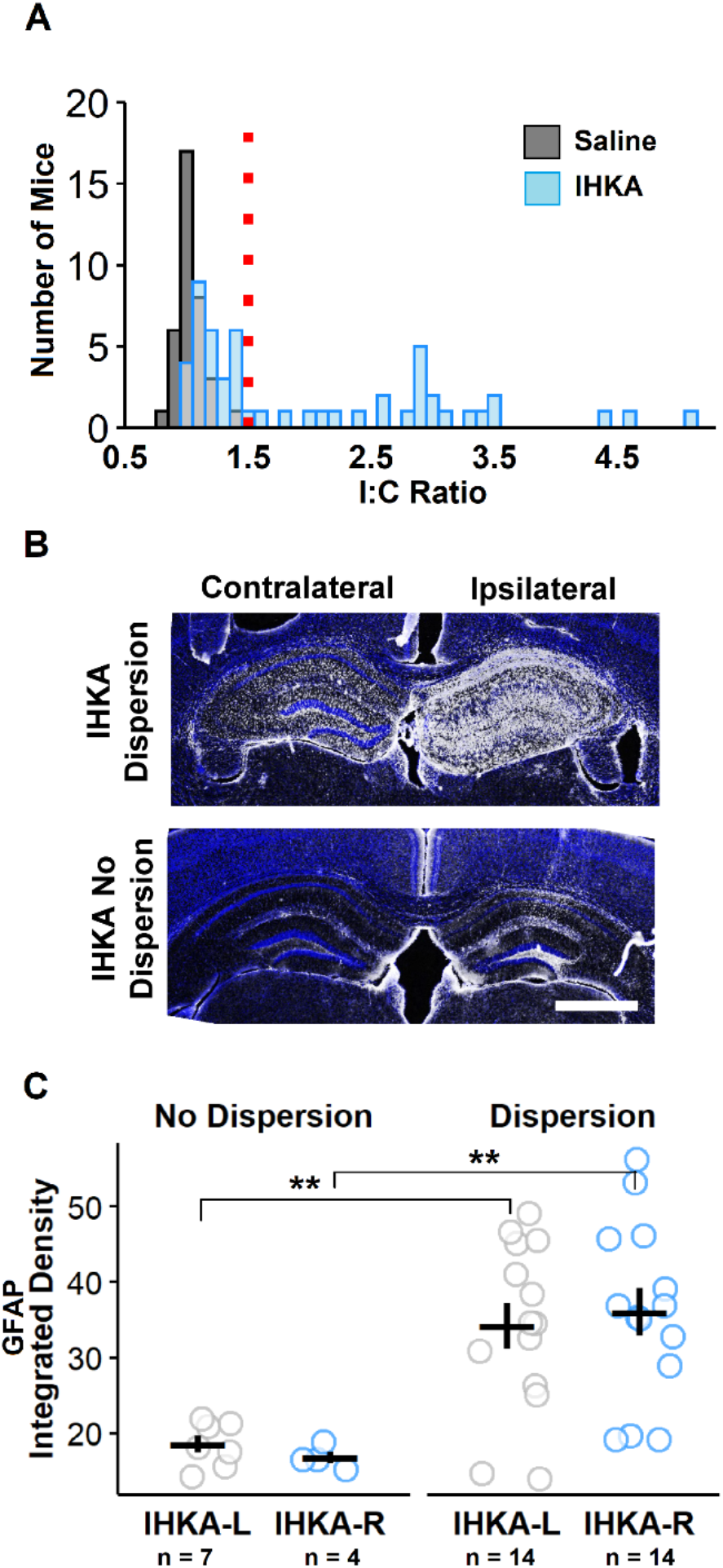
GFAP immunoreactivity does not differ based on side of IHKA injection. (A) Numbers of mice exhibiting I:C ratios. Red dotted line marks ratio value of of 1.5, which was selected as threshold to define animals that presented dispersion. (B) Representative images of GFAP staining. Scale bar = 1 mm. (B) Individual values and mean ± SEM of integrated density of GFAP staining normalized to overall size of ipsilateral hippocampus. **p < 0.01 for comparisions between injection side, and presence of dispersion by two way ANOVA and Tukey’s *post hoc* tests.

GFAP staining was evaluated in hippocampal sections of the same IHKA animals (availability of remaining tissue permitting) to determine whether similar differences in terms of gliosis based on injection site were also present (**Figure 3B**). Evaluation of GFAP intensity, normalized to ipsilateral hippocampal area, showed that, as expected, animals with dispersion showed higher levels of GFAP intensity than those without (F(1, 36) = 23.17, p = 2.66 x 10^-5^, **Figure 3C**), but there was no difference in the intensity of GFAP staining between IHKA-L and - R animals, regardless of whether dispersion was present or not (F(1, 36) = 0.08, p = 0.79, **Figure 3C**).

### 3.3 Estrous cycles are longer in IHKA mice

Estrous cycle monitoring was performed at 1, 2, and 4 months after saline or IHKA injection, and the cycle length was subsequently derived for each time point (**Figure 4A-B**). Cycle length was analyzed as it is the most robust measure of disruption in IHKA mice (Li et al., 2017, 2018, 2020). There was no effect of time after injection on the mean cycle length (F(2,419) = 1.15, p = 0.22), but there were differential shifts in this parameter across the months. At 1 month post-injection, longer cycles in comparison to respective control groups (F(1, 209) = 9.23, p = 0.002) were observed in both IHKA-L (p = 0.01) and IHKA-R (p = 0.003) mice, but this effect was not impacted by the side targeted for injection (F(1, 209) = 0.14, p = 0.70, **Figure 4C**). At 2 months post-injection, cycle length was still increased compared to controls (F(1,148) =6.61, p = 0.01) in IHKA-L (p = 0.05) and IHKA-R (p = 0.02) mice, again with no effects of injection site (F(1,148) = 0.49, p = 0.48, **Figure 4C**). At 4 months post-injection, only IHKA-R mice still exhibited increased cycle length compared to controls (p = 0.02, **Figure 4C**), but the cycle length at this time was not different between IHKA-R and IHKA-L groups (F(1,62) = 0.69, p = 0.41). Overall, mean cycle length was longer in all IHKA groups by 1 and 2 months post-injection, but at 4 months only the IHKA-R animals showed significantly lengthened cycles in comparison to controls. These results confirm previous findings of estrous cycle disruption occurring by two months after injection in IHKA-R mice (Li et al., 2017, 2018, 2020) and demonstrate that disruption can also be seen at one and four months post-injection in this group. Furthermore, these results extend previous findings by indicating that IHKA-L mice follow similar trends in cycle disruption as IHKA-R for the first 2 months after injection.

**Figure 4.**
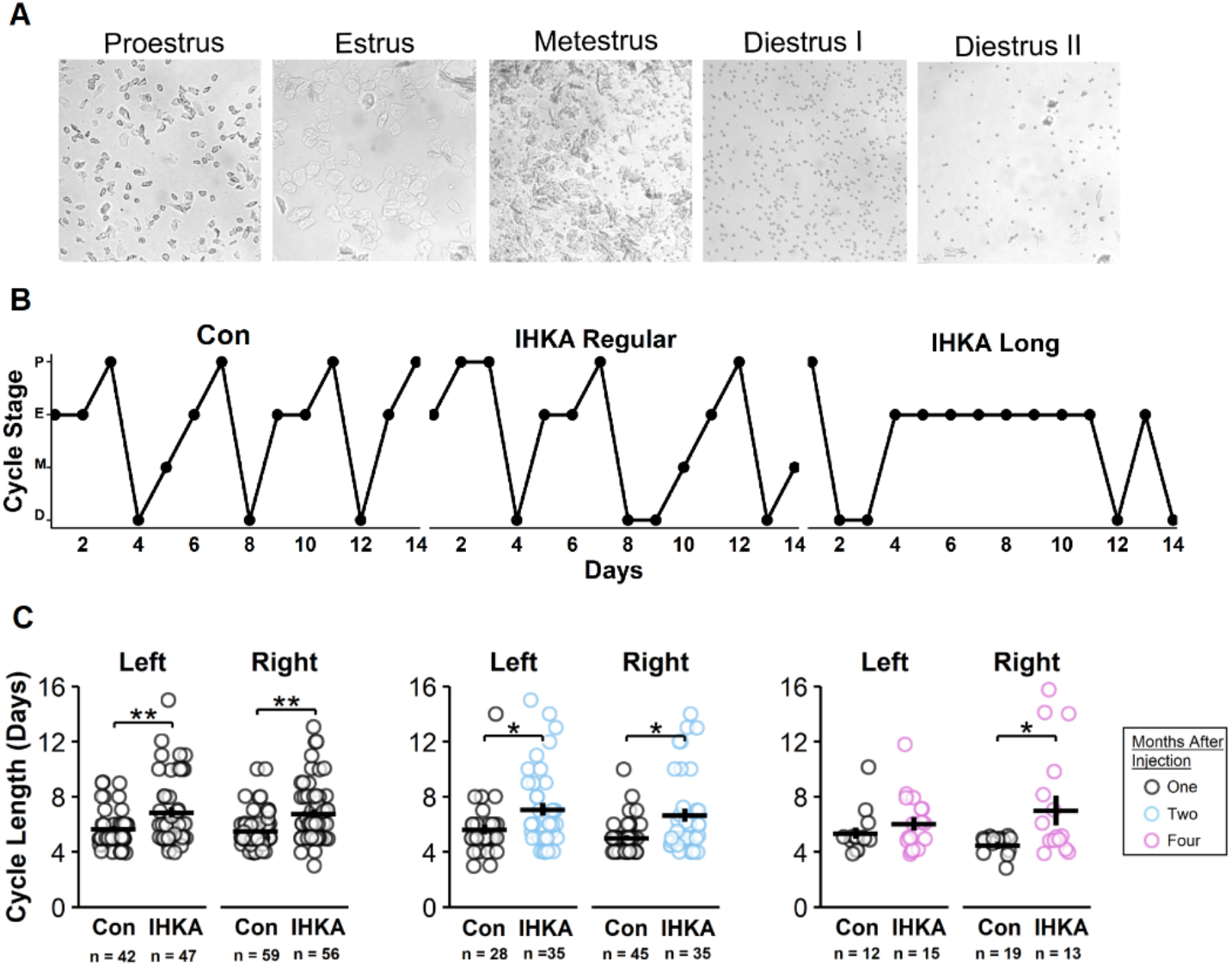
Estrous cycle period is lengthened after IHKA injection. (A) Vaginal cytology characteristics of each estrous cycle phase in the order of progression. The length of time to progress through each of these phases back to proestrus in the normal cycle thus takes 4-5 days. (B) Representative examples of a regular estrous cycle seen in a saline-injected control (left), an IHKA mouse with a regular 4-6 d cycle (middle), and an IHKA mouse with a long (≥7 day) cycle (right). (C) Cycle length of each mouse at each post-injection timepoint with mean ± SEM. * p < 0.05, ** p < 0.01 for comparisons between injection type (Con or IHKA), and injection side (left or right) and month timepoint (one, two, or four) by three-way ANOVA and Tukey’s *post hoc* tests.

The presence of IHKA mice with less granule cell dispersion (**Figure 2**), as well as the proportion of IHKA females that maintain a normal estrous cycle following injection (Li et al., 2017, 2018, 2020) (**Figure 4**), suggested that the degree of cycle disruption and the amount of dispersion may be correlated. However, there was no correlation between degree of granule cell dispersion and estrous cycle length (r^2^ = 0.03, p = 0.13).

### 3.4 Ovarian hormones and corticosterone are unchanged after IHKA injection

Changes in ovarian hormones such as estradiol, progesterone, and testosterone could potentially contribute to estrous cycle disruptions. In a previous study, decreased progesterone was observed at 2 months after injection in IHKA-R mice with long cycles (≥ 7-day cycle period), and IHKA-R mice with both long and regular cycles (4- to 6-day period) lacked the decrease in estradiol levels on estrus compared to diestrus seen in controls (Li et al., 2018). Furthermore, altered serum levels of these hormones have been reported in other rodent models of TLE that also display cycle disruption (Amado et al., 1993; Scharfman et al., 2008). Therefore, the trunk blood serum of IHKA-L, IHKA-R, Con-L, and Con-R mice collected on diestrus at 1, 2, and 4 months post-injection was analyzed through ELISA for levels of estradiol, progesterone, and testosterone. The time after injection did not have an effect on the concentrations of estradiol (F(2,96) = 0.56, p = 0.57), progesterone (F(2,105) = 0.56, p = 0.58), or testosterone (F(2,121) = 1.10, p = 0.33) (**Figure 5**). In addition, IHKA groups did not show differences in concentrations of estradiol (F(1,104) = 0.44, p = 0.50), progesterone (F(1,113) = 1.95, p=0.17), or testosterone (F(1,129) = 0.02, p=0.88) in comparison to controls. There were also no effects of injection side on the concentration of estradiol (F(1,104) = 0.22, p=0.63), progesterone (F(1,113) = 0.13, p=0.71), or testosterone (F(1,129) = 0.45, p =0.5). Furthermore, there was no correlation between progesterone levels and cycle length (r^2^ = 0.007, p = 0.52), in accordance with a previous study (Li et al., 2018). In addition, there was no significant effect of side of injection on the progesterone:estradiol ratio in each mouse, but there was a trend towards a difference between IHKA-R and saline-injected controls (F(1,77) = 2.91, p = 0.06).

**Figure 5.**
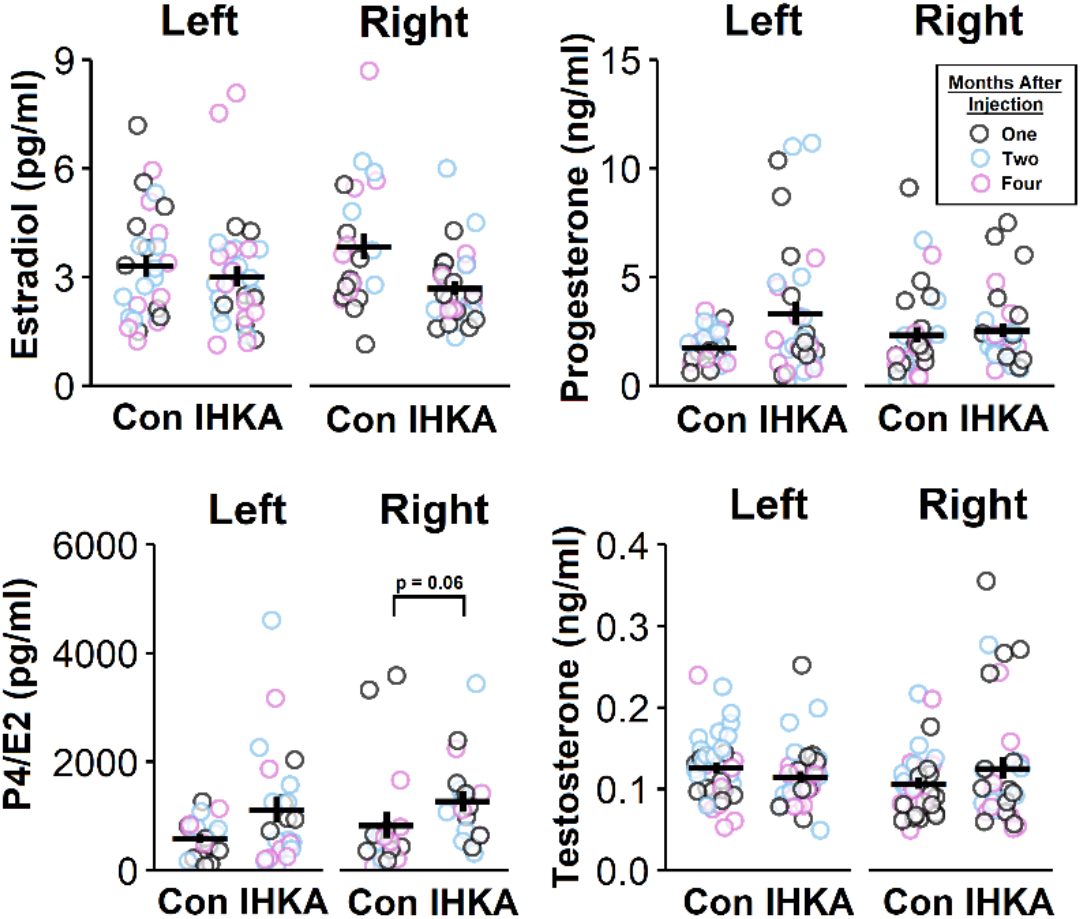
Ovarian hormones do not change with IHKA injection or laterality. Individual estradiol (top left), progesterone (top right), P4/E2 ratio (bottom left), and testosterone (bottom right) values for each mouse with mean ± SEM for each treatment. Comparisons between injection type (Con or IHKA), injection side (left or right), and ELISA plate made by three-way ANOVA.

High levels of the stress hormone corticosterone have been observed in animal models of epilepsy (Mazarati et al., 2009), potentially due to seizure-induced hyperexcitability of the hypothalamic-pituitary-adrenal axis (O’Toole et al., 2014). Increased corticosterone levels can drive increased seizure activity (Castro et al., 2012), and corticosterone can also disrupt the pulsatile pattern of pituitary LH release and subsequently alter the production of gonadal steroids (Kamel & Kubajak, 1987). Therefore, to evaluate the stress level of the animals between treatment groups and to determine whether stress was contributing to other hormonal findings, corticosterone was measured for the subset of animals for which sufficient serum remained after measuring the ovarian hormones. This evaluation showed that there was no significant effect of treatment (F(1,64) = 1.55, p = 0.21) or injection side (F(1,64) = 0.02, p = 0.87) on basal corticosterone concentrations (IHKA-R: 1122.03 ± 603.22 pg/ml, n = 23; IHKA-L: 1124.16 ± 637.30 pg/ml, n = 14; Con-R: 985.15 ± 460.98 pg/ml, n = 27; Con-L: 1020.11 ± 960.31 pg/ml, n = 5).

### 3.5 Numbers of ovarian follicles and corpora lutea are not changed after IHKA injection

The duration of time after injection had no effects on the number of primary (F(2,198) = 0.30, p = 0.74), secondary (F(2,198) = 2.90, p = 0.07), or antral (F(2,198) = 0.48, p = 0.62) follicles or corpora lutea (F(2,198) = 0.39, p = 0.68) in the ovaries. Therefore, the groups were collapsed across time of collection for further analysis. No significant differences in follicle or corpora lutea counts based on side of injection or saline vs. IHKA treatment were present (**Table 2**). Furthermore, no ovaries displayed cysts. These findings support and extend previous work that found no ovarian cysts or differences in follicle or corpora lutea counts in IHKA-R mice (Li et al., 2017).

**Table 2.** Ovarian follicle counts are unchanged with differing site of injection and independent of injection type. Means ± SEM of primary, secondary, and antral follicles as well as corpora lutea for each treatment group. Comparisons between injection type (Con or IHKA), and injection laterality (left or right) were made by two-way ANOVA.

|  | Control |  | IHKA |  | F-Value | P-Value |
| --- | --- | --- | --- | --- | --- | --- |
| Follicle Type | Left | Right | Left | Right |  |  |
| Primary | 1.66 ± 1.02 | 1.55 ± 1.17 | 1.51 ± 0.98 | 1.60 ± 1.07 | 0.22 | 0.88 |
| Secondary | 3.11 ± 1.80 | 2.80 ± 1.90 | 2.90 ± 1.73 | 3.21 ± 1.56 | 0.53 | 0.66 |
| Antral | 3.32 ± 1.27 | 3.49 ± 1.33 | 3.56 ± 1.47 | 3.57 ± 1.46 | 0.33 | 0.81 |
| Corpora Lutea | 2.27 ± 1.01 | 2.07 ± 1.07 | 1.84 ± 1.06 | 1.98 ± 1.06 | 1.34 | 0.26 |

### 3.6 Differences between IHKA-L and IHKA-R mice in pituitary expression of gonadotropin and Gnrhr genes

To evaluate the contribution of the pituitary to potential disruptions in reproductive function, *Fshb*, *Lhb,* and *Gnrhr* mRNA expression in the pituitaries of mice collected at 2 months after injection were measured using qPCR. *Fshb* expression was impacted by injection side (F(1,41) =6.87, p = 0.01), and type (F(1,41) = 5.88, p = 0.02) as levels were higher in IHKA-R than IHKA-L groups (p = 0.048), and in IHKA-R compared to Con-R (p = 0.004) (**Figure 6A**). *Lhb* expression was also impacted by injection side (F(1,41) = 10.19, p = 0.003) as IHKA-L mice showed higher expression of *Lhb* than IHKA-R (p = 0.01), but neither IHKA groups were significantly different from corresponding controls (F(1,41) = 4.35; left: p = 0.17; right: p = 0.99, **Figure 6B**). *Gnrhr* expression showed similar patterns as *Lhb,* with expression impacted by injection side (F(1,52) = 5.61, p = 0.02) with higher expression in IHKA-L than IHKA-R mice (p = 0.05), but neither group differed from corresponding saline controls (F(1,52) = 4.53; left: p = 0.1577; right: p = 0.68) **Figure 6C**). Overall, IHKA-R mice showed the highest elevation of *Fshb* expression, and IHKA-L showed higher *Lhb* and *Gnrhr* expression than IHKA-R mice.

**Figure 6.**
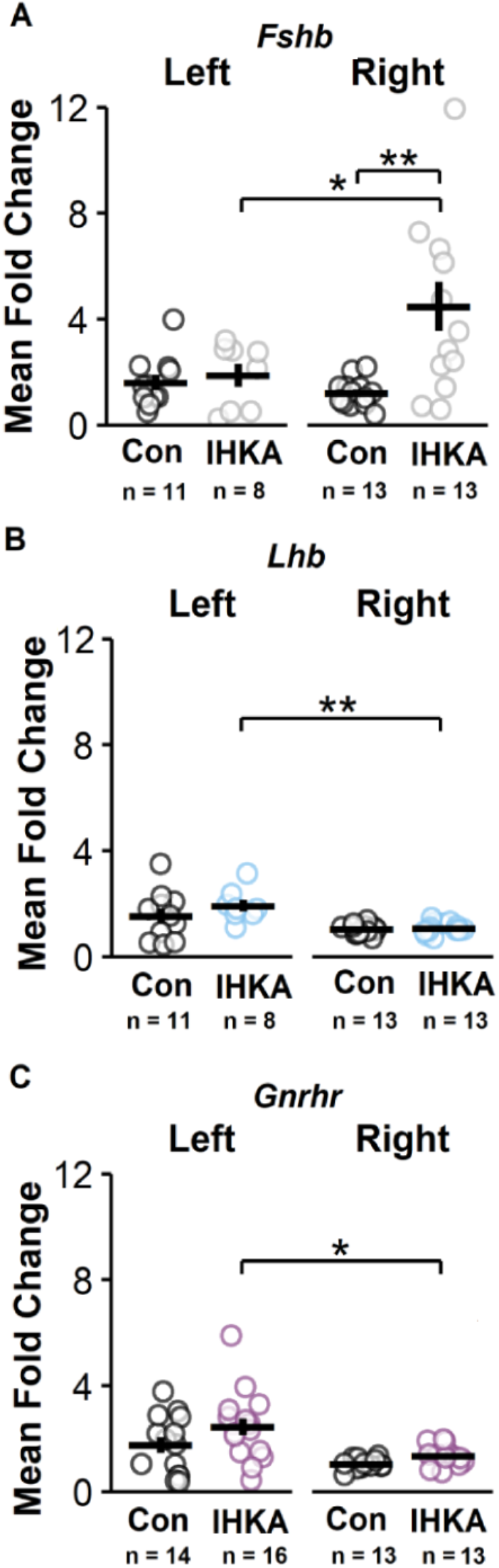
Lateralized differences in pituitary *Fshb, Lhb,* and *Gnrhr* gene expression two months after injection. Individual mean fold change values for control and IHKA mice for *Fshb* (A), *Lhb* (B), and *Gnrhr* (C), with mean ± SEM for each treatment group. * p ≤ 0.05, ** p < 0.01 for comparisons between injection type (Con or IHKA) and injection side (left or right) by two-way ANOVA and Tukey’s *post hoc* tests.

### 3.7 IHKA-L mice gain weight at an increased rate

In a previous study, IHKA-R mice were observed to have increased weight gain compared to controls (Li et al., 2017). To evaluate potential differences in weight gain based on side of injection, body weights were measured at the times of injection surgery and euthanasia. In comparison to Con-L mice, IHKA-L had higher weight gain at 1 month after injection (p = 0.0012) and borderline significantly higher weight gain at 2 months post-injection (p = 0.053) (**Figure 7**). However, the average weight gain in IHKA-L mice was not different based on time after injection. By contrast, IHKA-R mice did not display weight gain that differed from Con-R mice at any time after injection, but at 4 months after injection IHKA-R mice showed greater weight gain compared to those measured at 1 month (p = 0.0001). This result was similar to that observed in Con-L and -R groups, which also showed increased weight gain in mice measured at 2 months after injection compared to those measured at 1 month (Con-L: p = 0.02, Con-R: p = 0.002) as well as those mice measured at 2 months compared to those at 4 months (Con-L: p = 0.008, Con-R: p = 0.009). Overall, IHKA-L mice appeared to initially gain weight at a faster rate but then plateau at a stable weight, whereas IHKA-R mice showed a more gradual weight gain similar to that observed in controls over the same age range.

**Figure 7.**
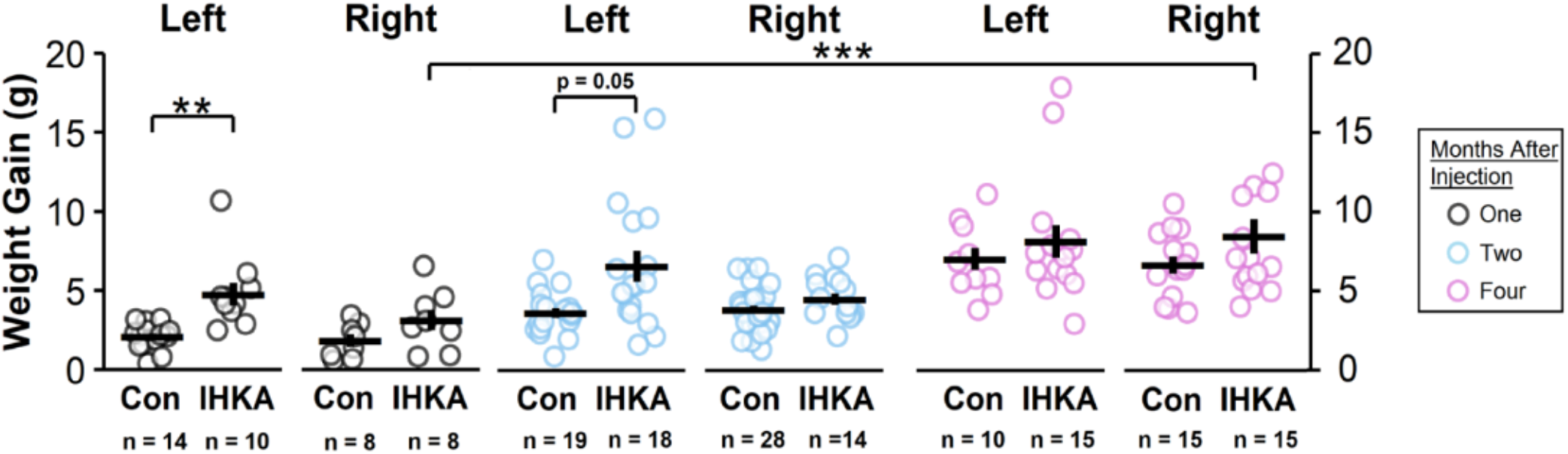
Post-injection weight gain of mice euthanized at one, two, or four months. Individual values and mean ± SEM for weight gain derived from differences in measurements at time of injection to time of tissue collection for each mouse. *p < 0.05, **p < 0.01 for comparisons between injection type (Con or IHKA), injection laterality (left or right), and post-injection time (one, two, or four months) by three-way ANOVA and Tukey’s *post hoc* tests.

## 4. Discussion

The findings in the present study indicate that targeting left or right dorsal hippocampus for KA injection has implications on the degree of hippocampal granule cell layer dispersion, phenotypes of pituitary gene expression, and the pattern of weight gain after injection in female C57BL/6J mice. However, there do not seem to be large phenotypic differences in hippocampal gliosis or in reproductive endocrine dysfunction presentation, as typified by increased estrous cycle length. The lack of differences in estrous cycle disruption based on the side of IHKA injection, as well as a lack of impact on ovarian morphology, supports and extends previous findings obtained in IHKA-R mice only (Li et al., 2017).

### 4.1 Lateralized differences in hippocampal granule cell dispersion in IHKA mice

The observed increase in granule cell layer dispersion and cell death in Ammon’s horn is commonly seen following IHKA injections (Bielefeld et al., 2017; Rattka et al., 2013) and a similar histological phenotype is often observed in patients with TLE (Berkovic et al., 1991). The results from this study further serve to highlight that this model recapitulates the variety of histopathological phenotypes seen in clinical TLE populations (Blümcke et al., 2013), as not all mice in the study with CA region cell death presented significant granule cell dispersion, and some IHKA mice did not exhibit dispersion or gliosis.

Furthermore, the present work provides a novel finding of a greater amount of dispersion present in IHKA-R compared to IHKA-L mice. Although less lateralized than the human brain, mice show structural and functional asymmetries in the hippocampus; wild-type C57BL/6J mice show greater granule cell layer volume in the right hippocampal dentate gyrus compared to the left (Tabibnia et al., 1999), and optogenetic silencing of the left CA3 region can impair long term spatial memory, but silencing the right CA3 does not produce the same effect (Shipton et al., 2014). Of note, a number of molecular, microanatomical, and behavioral asymmetries that have been reported in the rodent hippocampus (Jordan, 2020) could contribute to such differential presentation of this cellular dispersion. Intrinsic molecular asymmetries in mouse hippocampus, such as decreased expression of GluR1 in the cells of the left hippocampus compared to the right (Shinohara et al., 2008), could underlie distinct responses to KA, resulting in the differential presentation of dispersion in IHKA-L and -R animals. It should be noted, however, that the degree of gliosis was similar in the injected side of hippocampus in IHKA-L and -R animals, indicating a difference in lateralized phenotype between neuronal damage and astrocytic response. Higher severity of status epilepticus or chronic seizures that originate in the right temporal lobe versus the left could also contribute to differential death and reorganization of hippocampal cell populations. EEG recordings are needed to investigate the contributions of seizure severity to hippocampal neuron damage in IHKA-L and IHKA-R mice; such studies, beyond the scope of the present work, are currently ongoing in our lab.

### 4.2 Lateralized expression of gonadotropin and Gnrhr genes in pituitaries of IHKA mice

To understand whether changes at the level of the pituitary could at least partially underlie downstream reproductive endocrine disruption, mRNA expression levels of *Fshb*, *Lhb*, and *Gnrhr* were evaluated. The lack of differences in *Lhb* and *Gnrhr* levels in IHKA mice in comparison to controls, and only a mild increase in expression of each gene in IHKA-R compared to IHKA-L mice, suggest there might be an inherent difference in the pituitary’s response to the lateralization of damage inflicted by the injection, but that IHKA injection had no major effect on the transcription of these genes. By contrast, only IHKA-R mice showed increased expression of *Fshb*. *Fshb* synthesis is regulated not only by GnRH, but also by several other factors including inhibins, activins, and follistatins (Ying, 1988). *Lhb*, however, is mainly regulated by GnRH. As changes in inhibins, activins, and follistatins would only have effects on *Fshb*, these factors, rather than GnRH, may be more likely candidates in mediating this change. However, it should be noted that ovarian cytology was unchanged, and inhibins are produced from ovarian follicles and corpora lutea (Woodruff & Mayo, 1990). It is possible that locally produced activins (Corrigan et al., 1991) and follistatins (Ueno et al., 1987) differentially exert effects within the pituitary based on the side of hippocampus targeted for KA injection. Further evidence supporting a postulated role for these locally produced factors is the association of *Fshb* transcription with lower frequency of GnRH pulse input to the pituitary (Bernard & Brûlé, 2020). *Gnrhr* transcription is typically thought to be enhanced by increased amounts of GnRH (Cheon et al., 1999), and decreased in its absence. Therefore, if GnRH pulse frequency is decreased—as suggested by the *Fshb* transcript amounts—one might also expect a decrease in *Gnrhr* expression as well. However, this was not the case. As previously demonstrated in IHKA-R mice (Li et al., 2018), there may be altered release patterns of GnRH due to changes in the function of GnRH neurons and other hypothalamic afferents. The pituitary may then respond to this change by compensatorily maintaining GnRH-R expression so that activation is not subsequently decreased. The present findings are the first to demonstrate changes in pituitary gene expression in IHKA-injected animals. Furthermore, differences in expression profiles based on the side of IHKA injection indicate asymmetries in the seizure focus can have implications at the level of the pituitary.

### 4.3 Lengthening of estrous cycles in IHKA mice does not differ with side of injection

The lengthening of the average estrous cycle period is a phenotype that has been previously characterized in IHKA-R mice (Li et al., 2017, 2018, 2020) as well as in other rodent models of TLE (Amado et al., 1993; Edwards et al., 1999; Scharfman et al., 2008). Cycle disruption is noted here as a lengthening of the time it takes for a mouse to progress through each phase of the cycle. In a previous study, cycle disruption in IHKA-R mice did not develop until at least 6 weeks after injection (Li et al., 2017). In the present study, however, which incorporates a larger number of mice, estrous cycles were disrupted at one, two, and four months after injection. There may also be a more persistent effect in IHKA-R than in IHKA-L mice, potentially due to differential underlying mechanisms of the elongated cycles, as the degree of effect on cycle length diminished by 4 months in IHKA-L, but not IHKA-R, mice.

A lack of effect of the side of hippocampal injection on estrous cycle disruption was also seen in a previous study investigating the effects of left- vs. right-sided amygdala kindling in rats (Hum et al., 2009). This finding is consistent with the present results in IHKA mice, in which estrous cycle disruption was prominent but the side targeted for injection largely did not affect the overall presentation of the disruption. More prolonged estrous cycle disruption observed in IHKA-R than in IHKA-L mice might be linked to the asymmetries seen in hippocampal damage and changes in pituitary gene expression in these animals. However, estrous cycle disruption was otherwise largely phenotypically similar in both IHKA groups. Therefore, it could be that similar emergent reproductive endocrine phenotypes are produced from distinct underlying mechanisms at the neural and pituitary levels, thus masking the ability of IHKA or other mouse models to recapitulate the distinct reproductive endocrine outcomes seen in TLE patients with lateralized seizure foci.

### 4.4 Rate of post-injection weight gain in IHKA mice differs based on side targeted for injection

Incremental weight gain over time has been reported within the C57BL/6J mouse strain (Fahlström et al., 2011), which aligns with the patterns observed here in both saline-injected groups. The faster rate of weight gain in IHKA-L compared to Con-L mice is consistent with clinical findings of a potential correlation between left hippocampal atrophy and obesity in children (Mestre et al., 2017). It is also consistent with data from male rats indicating that orexigenic activation of the melanocortin system dominates on the left hypothalamic side (Kiss et al., 2020). In addition, female patients with TLE with left-sided seizure foci have a greater risk of developing PCOS (Herzog et al., 1986b), and women with PCOS typically have higher weight gain than the general population (Teede et al., 2013). The present findings suggest that the left hippocampus, and efferents to left hypothalamic regions, may have greater implications in weight regulation than the right hippocampus. It should be noted that the lack of difference in weight gain of IHKA-R mice in this study contrasts with previous findings of elevated body weight in IHKA-R female mice compared with controls (Li et al., 2017). Although the reported weight gain in the previous study for IHKA-R animals was higher than that observed in the present cohort of IHKA-R mice, it was similar to that of the IHKA-L animals in this study. A difference between the number of animals examined in each study, with a smaller cohort examined previously, as well as environmental factors including intermittent construction noise and vibration during the time of the present work, could have contributed to these discrepant results.

### 4.5 Ovarian hormones, follicles, and corpora lutea are unaffected by either IHKA or side of hippocampus targeted for injection

The lack of changes in the number of corpora lutea or follicles due to any of the treatment factors aligns with previous evaluations of ovarian morphology in IHKA-R mice (Li et al., 2017). In IHKA mice, no cysts were present in the ovary samples, and no changes in testosterone levels were observed. By contrast, systemic pilocarpine-treated female rats with epilepsy have shown prominent development of cysts and elevated testosterone (Scharfman et al., 2008); it remains unclear whether this difference represents a distinction based on chemoconvulsant mechanism of action and/or mode of injection, or a species difference between mice and rats. In this regard, C57BL/6 mice may be more resistant to age-related cyst development in comparison to other mouse strains (Kon et al., 2007), and mouse models of PCOS typically require powerful endocrine manipulations such as prenatal androgenization (Sullivan & Moenter, 2004) or chronic treatment with the aromatase inhibitor letrozole (Kauffman et al., 2015) to drive the presentation of cysts.

For consistency, collection of serum from trunk blood for gonadal hormone analysis was carried out on diestrus for all mice. As gonadal hormone levels fluctuate throughout the different estrous cycle stages, however, it is possible that changes in hormone levels between these groups could be occurring on other estrous cycle stages and are thus not represented in this data set. It should also be noted that previous findings suggested that IHKA mice with long cycles had altered progesterone levels correlated to cycle length (Li et al., 2018). This discrepancy could be attributed to tissue collections being performed at slightly different times of day in each study, or to mouse strain differences between the two studies (although the transgenic mice used in the previous study were on the C57BL/6J background). The previous study also used a population of mice bred in-house, whereas the mice in the present study were purchased and shipped to our facility at 6 weeks of age. Another factor could be that the ELISAs to measure hormone levels in the previous and present studies were performed in separate labs using different equipment. It should be noted, however, that similar mean and standard deviation values of progesterone levels were measured in both studies, suggesting that methodology is unlikely to be the prevailing source of variance. Corticosterone, which is often elevated with seizure activity (O’Toole et al., 2014; Wulsin et al., 2018; Zobel et al., 2004), has been known to disrupt the pulsatile release of LH and is linked with altered steroidogenesis at the ovary (Kamel & Kubajak, 1987). As corticosterone levels were unchanged in the animals in this study, however, there were no indications that physiological stress on the animals played a factor in the gonadal hormone results.

### 4.6 Concluding remarks

The present findings provide novel evidence of lateralized differences in phenotypic outcomes in female mice following IHKA injection, with significant differences in presentation of hippocampal granule cell dispersion, pituitary gonadotropin gene expression, and weight gain between IHKA-L and -R animals (**Figure 8**). The lateralized differences in hippocampal granule cell dispersion and pituitary gonadotropin gene expression, together with a lack of difference in estrous cycle lengthening phenotype based on injection site, indicate that there may be differential mechanisms that produce similar comorbid reproductive endocrine outcomes. This concept is akin to the recent postulate of latent sex differences in hippocampal function, in which the emergent output in males and females is similar but the underlying mechanisms are distinct (Jain et al., 2019; Koss et al., 2018). Whether this effect reflects compensation at the level of the pituitary, or effects further downstream in the hypothalamic-pituitary-ovarian axis, remains to be determined. Further analysis of pituitary gonadotropin release is needed to confirm if there are changes in functional FSH or LH levels that could promote cycle disruption in IHKA mice.

**Figure 8.**
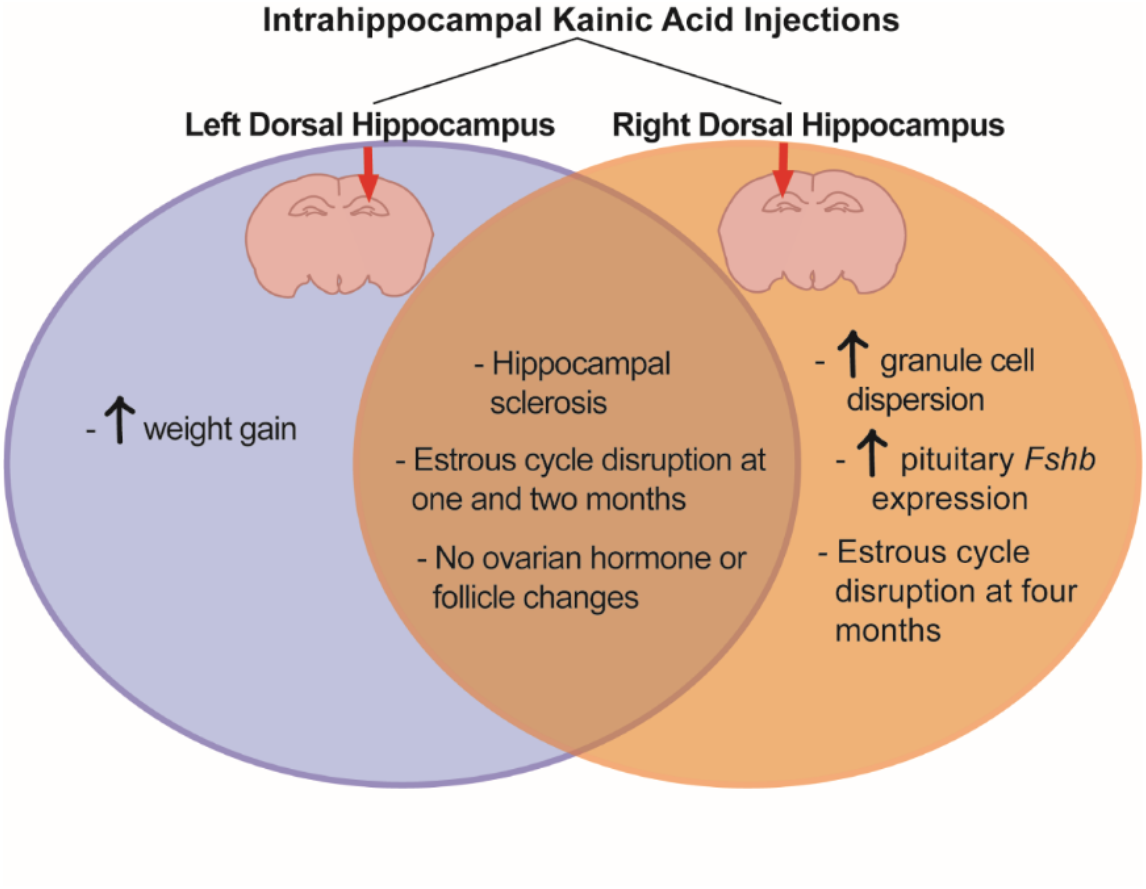
Summary of results. Outcomes of IHKA injections into the left or right dorsal hippocampus.

In summary, the present work demonstrates distinct lateralized differences in phenotypic outcome based on the side of the hippocampus targeted for IHKA injection, spanning from the level of the hippocampus to the pituitary to whole body weight. This work thus demonstrates the importance of choosing the appropriate hemisphere of hippocampus for initial epileptogenic insult and provides a foundation for future experimental design.

## Abbreviations

ELISA: enzyme-linked immunosorbent assay
FSH: follicle-stimulating hormone
GFAP: glial fibrillary acidic protein
GnRH: gonadotropin-releasing hormone
GnRH-R: gonadotropin-releasing hormone receptor
HPO: hypothalamic-pituitary-ovarian
IHKA: intrahippocampal kainic acid
KA: kainic acid
LH: luteinizing hormone
PCOS: polycystic ovary syndrome
TLE: temporal lobe epilepsy

## Acknowledgements

We thank Karen Doty for performing ovary sectioning and staining, Jiang Li for assistance with pilot studies for this project, and Manisha Naganatanahalli for assistance with preparing tissue for histology.

## Funding

This work was supported by the National Institutes of Health (NIH)/National Institute of Neurological Disorders and Stroke and the NIH Office of Research on Women’s Health through grants R03 NS103029 and R01 NS105825 (C.A.C.-H.).

## Author Contributions

C.A.C.-H. designed research; C.A.C., L.K.L., X.G., and R.Y. performed research; C.A.C., L.K.L., X.G., and L.T.R. analyzed data, C.A.C. and C.A.C.-H. wrote the paper.

## Notes

### Competing Interest Statement

The authors have declared no competing interest.

## References

Alessio, A., Bonilha, L., Rorden, C., Kobayashi, E., Min, L. L., Damasceno, B. P., & Cendes, F. (2006). Memory and language impairments and their relationships to hippocampal and perirhinal cortex damage in patients with medial temporal lobe epilepsy. Epilepsy & Behavior, 8(3), 593–600. https://doi.org/10.1016/j.yebeh.2006.01.007

Amado, D., Cavalheiro, E. A., & Bentivoglio, M. (1993). Epilepsy and hormonal regulation: The patterns of GnRH and galanin immunoreactivity in the hypothalamus of epileptic female rats. Epilepsy Research, 14(2), 149–159. https://doi.org/10.1016/0920-1211(93)90019-4

Berkovic, S. F., Andermann, F., Olivier, A., Ethier, R., Melanson, D., Robitaille, Y., Kuzniecky, R., Peters, T., & Feindel, W. (1991). Hippocampal sclerosis in temporal lobe epilepsy demonstrated by magnetic resonance imaging. Annals of Neurology, 29(2), 175–182. https://doi.org/10.1002/ana.410290210

Bernard, D. J., & Brûlé, E. (2020). Chapter 7 - Anterior Pituitary: Glycoprotein Hormones From Gonadotrope (FSH and LH) and Thyrotrope (TSH) Cells. In G. Litwack (Ed.), Hormonal Signaling in Biology and Medicine (pp. 119–144). Academic Press. https://doi.org/10.1016/B978-0-12-813814-4.00007-9

Bielefeld, P., Sierra, A., Encinas, J. M., Maletic-Savatic, M., Anderson, A., & Fitzsimons, C. P. (2017). A Standardized Protocol for Stereotaxic Intrahippocampal Administration of Kainic Acid Combined with Electroencephalographic Seizure Monitoring in Mice. Frontiers in Neuroscience, 11, 160. https://doi.org/10.3389/fnins.2017.00160

Blümcke, I., Thom, M., Aronica, E., Armstrong, D. D., Bartolomei, F., Bernasconi, A., Bernasconi, N., Bien, C. G., Cendes, F., Coras, R., Cross, J. H., Jacques, T. S., Kahane, P., Mathern, G. W., Miyata, H., Moshé, S. L., Oz, B., Özkara, Ç., Perucca, E., … Spreafico, R. (2013). International consensus classification of hippocampal sclerosis in temporal lobe epilepsy: A Task Force report from the ILAE Commission on Diagnostic Methods. Epilepsia, 54(7), 1315–1329. https://doi.org/10.1111/epi.12220

Blümcke, I., Thom, M., & Wiestler, O. D. (2002). Ammon’s horn sclerosis: A maldevelopmental disorder associated with temporal lobe epilepsy. Brain Pathology (Zurich, Switzerland), 12(2), 199–211.

Bonilha, L., Rorden, C., Halford, J. J., Eckert, M., Appenzeller, S., Cendes, F., & Li, L. M. (2007). Asymmetrical extra-hippocampal grey matter loss related to hippocampal atrophy in patients with medial temporal lobe epilepsy. Journal of Neurology, Neurosurgery & Psychiatry, 78(3), 286–294. https://doi.org/10.1136/jnnp.2006.103994

Bouilleret, V., Ridoux, V., Depaulis, A., Marescaux, C., Nehlig, A., & Le Gal La Salle, G. (1999). Recurrent seizures and hippocampal sclerosis following intrahippocampal kainate injection in adult mice: Electroencephalography, histopathology and synaptic reorganization similar to mesial temporal lobe epilepsy. Neuroscience, 89(3), 717–729. https://doi.org/10.1016/s0306-4522(98)00401-1

Campos, B. M. de, Coan, A. C., Yasuda, C. L., Casseb, R. F., & Cendes, F. (2016). Large-scale brain networks are distinctly affected in right and left mesial temporal lobe epilepsy. Human Brain Mapping, 37(9), 3137–3152. https://doi.org/10.1002/hbm.23231

Castro, O. W., Santos, V. R., Pun, R. Y. K., McKlveen, J. M., Batie, M., Holland, K. D., Gardner, M., Garcia-Cairasco, N., Herman, J. P., & Danzer, S. C. (2012). Impact of corticosterone treatment on spontaneous seizure frequency and epileptiform activity in mice with chronic epilepsy. PloS One, 7(9), e46044. https://doi.org/10.1371/journal.pone.0046044

Cheon, M., Park, D., Kim, K., Park, S. D., & Ryu, K. (1999). Homologous upregulation of GnRH receptor mRNA by continuous GnRH in cultured rat pituitary cells. Endocrine, 11(1), 49– 55. https://doi.org/10.1385/ENDO:11:1:49

Christian, C. A., Reddy, D. S., Maguire, J., & Forcelli, P. A. (2020). Sex Differences in the Epilepsies and Associated Comorbidities: Implications for Use and Development of Pharmacotherapies. Pharmacological Reviews, 72(4), 767–800. https://doi.org/10.1124/pr.119.017392

Coan, A. C., Appenzeller, S., Bonilha, L., Li, L. M., & Cendes, F. (2009). Seizure frequency and lateralization affect progression of atrophy in temporal lobe epilepsy. Neurology, 73(11), 834–842. https://doi.org/10.1212/WNL.0b013e3181b783dd

Corrigan, A. Z., Bilezikjian, L. M., Carroll, R. S., Bald, L. N., Schmelzer, C. H., Fendly, B. M., Mason, A. J., Chin, W. W., Schwall, R. H., & Vale, W. (1991). Evidence for an autocrine role of activin B within rat anterior pituitary cultures. Endocrinology, 128(3), 1682–1684. https://doi.org/10.1210/endo-128-3-1682

Edwards, H. E., Burnham, W. M., Ng, M. M., Asa, S., & MacLusky, N. J. (1999). Limbic seizures alter reproductive function in the female rat. Epilepsia, 40(10), 1370–1377. https://doi.org/10.1111/j.1528-1157.1999.tb02007.x

Engel, J. (2001). Mesial temporal lobe epilepsy: What have we learned? The Neuroscientist: A Review Journal Bringing Neurobiology, Neurology and Psychiatry, 7(4), 340–352. https://doi.org/10.1177/107385840100700410

Fahlström, A., Yu, Q., & Ulfhake, B. (2011). Behavioral changes in aging female C57BL/6 mice. Neurobiology of Aging, 32(10), 1868–1880. https://doi.org/10.1016/j.neurobiolaging.2009.11.003

Gazzaniga, M. S. (1995). Principles of human brain organization derived from split-brain studies. Neuron, 14(2), 217–228. https://doi.org/10.1016/0896-6273(95)90280-5

Gross, D. W., Concha, L., & Beaulieu, C. (2006). Extratemporal White Matter Abnormalities in Mesial Temporal Lobe Epilepsy Demonstrated with Diffusion Tensor Imaging. Epilepsia, 47(8), 1360–1363. https://doi.org/10.1111/j.1528-1167.2006.00603.x

Gröticke, I., Hoffmann, K., & Löscher, W. (2008). Behavioral alterations in a mouse model of temporal lobe epilepsy induced by intrahippocampal injection of kainate. Experimental Neurology, 213(1), 71–83. https://doi.org/10.1016/j.expneurol.2008.04.036

Haneef, Z., Lenartowicz, A., Yeh, H. J., Levin, H. S., Engel, J., & Stern, J. M. (2014). Functional connectivity of hippocampal networks in temporal lobe epilepsy. Epilepsia, 55(1), 137– 145. https://doi.org/10.1111/epi.12476

Helmstaedter, C., & Kockelmann, E. (2006). Cognitive Outcomes in Patients with Chronic Temporal Lobe Epilepsy. Epilepsia, 47(s2), 96–98. https://doi.org/10.1111/j.1528-1167.2006.00702.x

Herzog, A. G. (1993). A relationship between particular reproductive endocrine disorders and the laterality of epileptiform discharges in women with epilepsy. Neurology, 43(10), 1907–1910. https://doi.org/10.1212/wnl.43.10.1907

Herzog, A. G., Seibel, M. M., Schomer, D. L., Vaitukaitis, J. L., & Geschwind, N. (1986a). Reproductive endocrine disorders in men with partial seizures of temporal lobe origin. Archives of Neurology, 43(4), 347–350. https://doi.org/10.1001/archneur.1986.00520040035015

Herzog, A. G., Seibel, M. M., Schomer, D. L., Vaitukaitis, J. L., & Geschwind, N. (1986b). Reproductive endocrine disorders in women with partial seizures of temporal lobe origin. Archives of Neurology, 43(4), 341–346. https://doi.org/10.1001/archneur.1986.00520040029014

Hum, K. M., Megna, S., & Burnham, W. M. (2009). The effects of right and left amygdala kindling on the female reproductive system in rats. Epilepsia, 50(4), 880–886. https://doi.org/10.1111/j.1528-1167.2008.01982.x

Jain, A., Huang, G. Z., & Woolley, C. S. (2019). Latent Sex Differences in Molecular Signaling That Underlies Excitatory Synaptic Potentiation in the Hippocampus. Journal of Neuroscience, 39(9), 1552–1565. https://doi.org/10.1523/JNEUROSCI.1897-18.2018

Jordan, J. T. (2020). The rodent hippocampus as a bilateral structure: A review of hemispheric lateralization. Hippocampus, 30(3), 278–292. https://doi.org/10.1002/hipo.23188

Kalinin, V. V., & Zheleznova, E. V. (2007). Chronology and evolution of temporal lobe epilepsy and endocrine reproductive dysfunction in women: Relationships to side of focus and catameniality. Epilepsy & Behavior: E&B, 11(2), 185–191. https://doi.org/10.1016/j.yebeh.2007.04.014

Kamel, F., & Kubajak, C. L. (1987). Modulation of gonadotropin secretion by corticosterone: Interaction with gonadal steroids and mechanism of action. Endocrinology, 121(2), 561– 568. https://doi.org/10.1210/endo-121-2-561

Kandratavicius, L., Lopes-Aguiar, C., Bueno-Júnior, L. S., Romcy-Pereira, R. N., Hallak, J. E. C., & Leite, J. P. (2012). Psychiatric comorbidities in temporal lobe epilepsy: Possible relationships between psychotic disorders and involvement of limbic circuits. Revista Brasileira De Psiquiatria (Sao Paulo, Brazil: 1999), 34(4), 454–466. https://doi.org/10.1016/j.rbp.2012.04.007

Kauffman, A. S., Thackray, V. G., Ryan, G. E., Tolson, K. P., Glidewell-Kenney, C. A., Semaan, S. J., Poling, M. C., Iwata, N., Breen, K. M., Duleba, A. J., Stener-Victorin, E., Shimasaki, S., Webster, N. J., & Mellon, P. L. (2015). A Novel Letrozole Model Recapitulates Both the Reproductive and Metabolic Phenotypes of Polycystic Ovary Syndrome in Female Mice. Biology of Reproduction, 93(3), 69. https://doi.org/10.1095/biolreprod.115.131631

Keller, S. S., & Roberts, N. (2008). Voxel-based morphometry of temporal lobe epilepsy: An introduction and review of the literature. Epilepsia, 49(5), 741–757. https://doi.org/10.1111/j.1528-1167.2007.01485.x

Keller, S. S., Schoene-Bake, J.-C., Gerdes, J. S., Weber, B., & Deppe, M. (2012). Concomitant Fractional Anisotropy and Volumetric Abnormalities in Temporal Lobe Epilepsy: Cross-Sectional Evidence for Progressive Neurologic Injury. PLOS ONE, 7(10), e46791. https://doi.org/10.1371/journal.pone.0046791

Kiss, D. S., Toth, I., Jocsak, G., Bartha, T., Frenyo, L. V., Barany, Z., Horvath, T. L., & Zsarnovszky, A. (2020). Metabolic Lateralization in the Hypothalamus of Male Rats Related to Reproductive and Satiety States. Reproductive Sciences (Thousand Oaks, Calif.), 27(5), 1197–1205. https://doi.org/10.1007/s43032-019-00131-3

Kon, Y., Konno, A., Hashimoto, Y., & Endoh, D. (2007). Ovarian cysts in MRL/MpJ mice originate from rete ovarii. Anatomia, Histologia, Embryologia, 36(3), 172–178. https://doi.org/10.1111/j.1439-0264.2006.00728.x

Koss, W. A., Haertel, J. M., Philippi, S. M., & Frick, K. M. (2018). Sex Differences in the Rapid Cell Signaling Mechanisms Underlying the Memory-Enhancing Effects of 17β-Estradiol. ENeuro, 5(5), ENEURO.0267-18.2018. https://doi.org/10.1523/ENEURO.0267-18.2018

Li, J., Kim, J. S., Abejuela, V. A., Lamano, J. B., Klein, N. J., & Christian, C. A. (2017). Disrupted female estrous cyclicity in the intrahippocampal kainic acid mouse model of temporal lobe epilepsy. Epilepsia Open, 2(1), 39–47. https://doi.org/10.1002/epi4.12026

Li, J., Leverton, L. K., Naganatanahalli, L. M., & Christian-Hinman, C. A. (2020). Seizure burden fluctuates with the female reproductive cycle in a mouse model of chronic temporal lobe epilepsy. Experimental Neurology, 334, 113492. https://doi.org/10.1016/j.expneurol.2020.113492

Li, J., Robare, J. A., Gao, L., Ghane, M. A., Flaws, J. A., Nelson, M. E., & Christian, C. A. (2018). Dynamic and Sex-Specific Changes in Gonadotropin-Releasing Hormone Neuron Activity and Excitability in a Mouse Model of Temporal Lobe Epilepsy. ENeuro, 5(5), ENEURO.0273-18.2018. https://doi.org/10.1523/ENEURO.0273-18.2018

Lisgaras, C. P., & Scharfman, H. E. (2021). Robust chronic convulsive seizures, high frequency oscillations, and human seizure onset patterns in an intrahippocampal kainic acid model in mice BioRxiv (p. 2021.06.28.450253). https://doi.org/10.1101/2021.06.28.450253

Mazarati, A. M., Shin, D., Kwon, Y. S., Bragin, A., Pineda, E., Tio, D., Taylor, A. N., & Sankar, R. (2009). Elevated plasma corticosterone level and depressive behavior in experimental temporal lobe epilepsy. Neurobiology of Disease, 34(3), 457–461. https://doi.org/10.1016/j.nbd.2009.02.018

Mestre, Z. L., Bischoff-Grethe, A., Eichen, D. M., Wierenga, C. E., Strong, D., & Boutelle, K. N. (2017). Hippocampal atrophy and altered brain responses to pleasant tastes among obese compared with healthy weight children. International Journal of Obesity (2005), 41(10), 1496–1502. https://doi.org/10.1038/ijo.2017.130

Myers, M., Britt, K. L., Wreford, N. G. M., Ebling, F. J. P., & Kerr, J. B. (2004). Methods for quantifying follicular numbers within the mouse ovary. Reproduction (Cambridge, England), 127(5), 569–580. https://doi.org/10.1530/rep.1.00095

Nantie, L. B., Himes, A. D., Getz, D. R., & Raetzman, L. T. (2014). Notch signaling in postnatal pituitary expansion: Proliferation, progenitors, and cell specification. Molecular Endocrinology (Baltimore, Md.), 28(5), 731–744. https://doi.org/10.1210/me.2013-1425

O’Toole, K. K., Hooper, A., Wakefield, S., & Maguire, J. (2014). Seizure-induced disinhibition of the HPA axis increases seizure susceptibility. Epilepsy Research, 108(1), 29–43. https://doi.org/10.1016/j.eplepsyres.2013.10.013

Pantier, L. K., Li, J., & Christian, C. A. (2019). Estrous Cycle Monitoring in Mice with Rapid Data Visualization and Analysis. Bio-Protocol, 9(17), e3354. https://doi.org/10.21769/BioProtoc.3354

Paxinos, G., & Keith B.J. Franklin. (2019). The Mouse Brain in Stereotaxic Coordinates (5th ed.). Academic Press.

Phuong, T. H., Houot, M., Méré, M., Denos, M., Samson, S., & Dupont, S. (2021). Cognitive impairment in temporal lobe epilepsy: Contributions of lesion, localization and lateralization. Journal of Neurology, 268(4), 1443–1452. https://doi.org/10.1007/s00415-020-10307-6

Racine, R. J. (1972). Modification of seizure activity by electrical stimulation. II. Motor seizure. Electroencephalography and Clinical Neurophysiology, 32(3), 281–294. https://doi.org/10.1016/0013-4694(72)90177-0

Rattka, M., Brandt, C., & Löscher, W. (2013). The intrahippocampal kainate model of temporal lobe epilepsy revisited: Epileptogenesis, behavioral and cognitive alterations, pharmacological response, and hippoccampal damage in epileptic rats. Epilepsy Research, 103(2–3), 135–152. https://doi.org/10.1016/j.eplepsyres.2012.09.015

Riban, V., Bouilleret, V., Pham-Lê, B. T., Fritschy, J.-M., Marescaux, C., & Depaulis, A. (2002). Evolution of hippocampal epileptic activity during the development of hippocampal sclerosis in a mouse model of temporal lobe epilepsy. Neuroscience, 112(1), 101–111. https://doi.org/10.1016/s0306-4522(02)00064-7

Scharfman, H. E., Kim, M., Hintz, T. M., & MacLusky, N. J. (2008). Seizures and reproductive function: Insights from female rats with epilepsy. Annals of Neurology, 64(6), 687–697. https://doi.org/10.1002/ana.21518

Shinohara, Y., Hirase, H., Watanabe, M., Itakura, M., Takahashi, M., & Shigemoto, R. (2008). Left-right asymmetry of the hippocampal synapses with differential subunit allocation of glutamate receptors. Proceedings of the National Academy of Sciences of the United States of America, 105(49), 19498–19503. https://doi.org/10.1073/pnas.0807461105

Shipton, O. A., El-Gaby, M., Apergis-Schoute, J., Deisseroth, K., Bannerman, D. M., Paulsen, O., & Kohl, M. M. (2014). Left–right dissociation of hippocampal memory processes in mice. Proceedings of the National Academy of Sciences, 111(42), 15238–15243. https://doi.org/10.1073/pnas.1405648111

Sullivan, S. D., & Moenter, S. M. (2004). Prenatal androgens alter GABAergic drive to gonadotropin-releasing hormone neurons: Implications for a common fertility disorder. Proceedings of the National Academy of Sciences of the United States of America, 101(18), 7129–7134. https://doi.org/10.1073/pnas.0308058101

Sutula, T., Cascino, G., Cavazos, J., Parada, I., & Ramirez, L. (1989). Mossy fiber synaptic reorganization in the epileptic human temporal lobe. Annals of Neurology, 26(3), 321– 330. https://doi.org/10.1002/ana.410260303

Tabibnia, G., Cooke, B. M., & Breedlove, S. M. (1999). Sex difference and laterality in the volume of mouse dentate gyrus granule cell layer. Brain Research, 827(1–2), 41–45. https://doi.org/10.1016/s0006-8993(99)01262-7

Teede, H. J., Joham, A. E., Paul, E., Moran, L. J., Loxton, D., Jolley, D., & Lombard, C. (2013). Longitudinal weight gain in women identified with polycystic ovary syndrome: Results of an observational study in young women. Obesity (Silver Spring, Md.), 21(8), 1526–1532. https://doi.org/10.1002/oby.20213

Ueno, N., Ling, N., Ying, S. Y., Esch, F., Shimasaki, S., & Guillemin, R. (1987). Isolation and partial characterization of follistatin: A single-chain Mr 35,000 monomeric protein that inhibits the release of follicle-stimulating hormone. Proceedings of the National Academy of Sciences of the United States of America, 84(23), 8282–8286. https://doi.org/10.1073/pnas.84.23.8282

Woodruff, T. K., & Mayo, K. E. (1990). Regulation of inhibin synthesis in the rat ovary. Annual Review of Physiology, 52, 807–821. https://doi.org/10.1146/annurev.ph.52.030190.004111

Wulsin, A. C., Franco-Villanueva, A., Romancheck, C., Morano, R. L., Smith, B. L., Packard, B. A., Danzer, S. C., & Herman, J. P. (2018). Functional disruption of stress modulatory circuits in a model of temporal lobe epilepsy. PloS One, 13(5), e0197955. https://doi.org/10.1371/journal.pone.0197955

Ying, S. Y. (1988). Inhibins, activins, and follistatins: Gonadal proteins modulating the secretion of follicle-stimulating hormone. Endocrine Reviews, 9(2), 267–293. https://doi.org/10.1210/edrv-9-2-267

Zobel, A., Wellmer, J., Schulze-Rauschenbach, S., Pfeiffer, U., Schnell, S., Elger, C., & Maier, W. (2004). Impairment of inhibitory control of the hypothalamic pituitary adrenocortical system in epilepsy. European Archives of Psychiatry and Clinical Neuroscience, 254(5), 303–311. https://doi.org/10.1007/s00406-004-0499-9

